# Coordinated development of the male and female reproductive systems in the cestode *Hymenolepis microstoma*

**DOI:** 10.64898/2026.06.15.732408

**Authors:** Emilia Failache, Matías Preza, Jimena Montagne, Marc Kaethner, Uriel Koziol

## Abstract

**Background:** Cestodes have complex hermaphroditic reproductive systems that produce massive numbers of eggs. This reproductive output is made possible by the continuous production of serially repeated sets of reproductive systems (proglottids). However, their reproductive development remains poorly understood.

**Results:** We characterized reproductive development in the model cestode *Hymenolepis microstoma* by analyzing markers of cell proliferation, meiosis, and differentiation along the series of proglottids. Reproductive development begins with the formation of a central genital primordium, from which the reproductive ducts and gonads differentiate. Development is proterandrous, and testicular development is prolonged. In contrast, female reproductive development occurs over a short interval and is characterized by the coordinated differentiation of the ovary and vitelline gland. Entry of oocytes into meiosis is almost synchronous, and paralleled by cell proliferation in the vitelline gland. Subsequent growth of arrested oocytes and differentiation of vitelline cells occur in parallel. Insemination coincides with the onset of ovarian meiosis, indicating a close temporal coordination between male and female reproductive development. Finally, we show that gametogenesis and insemination proceed in adult worms maintained *in vitro*.

**Conclusions:** Our findings show the coordination of reproductive development in a self-fertile hermaphrodite, and provide an experimental system for studying reproductive development in cestodes.

## 1. Introduction

Parasitic flatworms, including cestodes, trematodes, and monogeneans, exhibit some of the most remarkable reproductive capacities among animals. Their massive reproductive output is thought to be essential for maintaining complex life cycles involving transmission between hosts. Among parasitic flatworms, cestodes display body segmentation (proglottization), a major evolutionary innovation in their reproductive development.^1^ Whereas a few basal cestode groups are monozoic, most cestodes possess a segmented body, the strobila, composed of serially repeated units termed proglottids, each containing an independent hermaphroditic reproductive system.^1–3^ New proglottids are generated continuously from the neck, a germinative region located posterior to the scolex (head). This region contains numerous proliferative undifferentiated cells, known as germinative cells in cestodes, which are morphologically and functionally similar to the neoblasts (stem cells) of free-living flatworms such as planarians.^4–8^ Since proglottids are formed successively from the neck, newly generated proglottids are located near this region, whereas progressively older, more developed proglottids are found toward the posterior end of the strobila.

During the development of each new proglottid, a genital primordium is formed as a localized accumulation of germinative cells.^5,9–11^ The male and female reproductive systems differentiate from this primordium, including the reproductive ducts, the gonads (testes and ovaries), and their associated glands, including the vitelline glands. Vitelline glands are a characteristic feature of neoophoran flatworms, a clade including parasitic flatworms together with several free-living groups.^2,12^ Unlike other animals in which yolk is incorporated into the oocyte, neoophoran flatworms possess alecithal oocytes and produce specialized cells in the vitelline gland (vitellocytes), which contribute eggshell material and nutritional reserves to developing embryos.^2,13,14^ In derived groups of cestodes such as in the order Cyclophyllidea, vitellocytes contain small or negligible nutritional reserves, and nutrition depends largely on resources provided within the uterus by the progenitor.^14^

Because proglottids are continuously produced, and reproductive development proceeds in each proglottid in an ordered sequence, the strobila constitutes a developmental series in which reproductive structures are found at successive stages of differentiation. Immature proglottids (located anteriorly) are followed by mature proglottids containing fully differentiated reproductive systems and subsequently by gravid proglottids filled with eggs (located more posteriorly). Spermatogenesis in cestodes has been extensively studied, particularly as sperm ultrastructure has been used as a source of useful phylogenetic characters.^15–19^ Spermatogonia undergo incomplete cell divisions, producing characteristic rosettes of two, four, and eight interconnected cells that divide synchronously. These generate 16 primary spermatocytes, which undergo two meiotic divisions to generate clusters of 64 spermatids, finally differentiating into mature spermatozoa. Female gonadal development generally occurs later (proterandry), and their development has only been studied in any detail in a handful of cestode species.^16,20–22^ Oocytes differentiate from oogonia within the ovary and are released while arrested in prophase I of meiosis. Fertilization occurs after their release, and the fertilized oocyte subsequently completes both meiotic divisions.^1,15^ Zygotes associate with vitellocytes within the oviduct, leading to egg formation as the vitellocytes secrete their eggshell globules.^14^ Eggs accumulate within the uterus and, in many species, gravid proglottids are eventually shed through apolysis, allowing their release from the definitive host into the external environment.

Among cestodes, species of *Hymenolepis* have been classical experimental models because of the simple maintenance of their life cycle under laboratory conditions and their phylogenetic proximity to other cyclophyllidean species of medical and veterinary relevance.^23,24^ Recent advances have expanded the experimental toolkit available for these organisms.^7,25–30^ Nevertheless, despite the importance of reproductive development in their life cycle, fundamental aspects of this process remain poorly understood in cestodes generally and in *Hymenolepis* in particular. Little is known regarding the sequence of events underlying the differentiation of male and female reproductive systems, the coordination between their developmental programs, and the coordination between ovarian and vitelline development within the female system. This coordination is particularly important because, unlike in many other animals, the gonads of individual proglottids do not continuously produce gametes for extended periods of time. Instead, gametogenesis, fertilization, and egg production appear to occur during a relatively restricted window, after which gonadal tissues degenerate as the uterus expands.

Classical histological studies of reproductive development in the rat tapeworm *Hymenolepis diminuta* described the early formation of a central genital primordium, from which a lateral extension gives rise to the male and female reproductive ducts.^11^ Testes subsequently develop laterally, while the ovary and vitelline gland derive from the central regions of the primordium. Spermatogenesis in this species was also analyzed at the ultrastructural level and is typical for cestodes.^31,32^ Pulse-chase assays with tritiated thymidine *in vivo* have shown a rapid progression through the different stages of spermatogenesis (requiring fewer than 48 hours from primary spermatocytes to mature sperm^33^). Many details of the reproductive anatomy of *H. diminuta* could also be revealed using fluorescent lectins and antibodies for widely conserved epitopes.^26^ Reproductive development in the mouse tapeworm, *Hymenolepis microstoma*, has not been investigated in detail, but is thought to be similar. *H. microstoma* is a self-fertile hermaphrodite.^34^ The development from genital primordia to fully mature proglottids requires approximately one week, and the mature proglottids only last for two days before the ovary disappears as the uterus becomes filled with eggs.^35^ More recently, Olson et al. (2018)^27^ have analyzed the differential gene expression patterns of different regions of the strobila of *H. microstoma*, identifying several genes specifically expressed in the developing gonads and vitelline gland.

In mature proglottids of *Hymenolepis* spp.^16,23^, the male reproductive system consists of three testes connected by efferent ducts to a vas deferens, which terminates in an eversible copulatory organ, the cirrus, at the right lateral margin of each proglottid. The vas deferens enlarges to form two successive seminal vesicles (one external and one internal to the muscular cirrus pouch), and is associated with unicellular prostatic glands. The female reproductive system comprises a centrally located ovary and a posterior vitelline gland. The vagina opens into a common genital atrium shared with the cirrus and leads to a seminal receptacle that stores spermatozoa. The seminal receptacle communicates with the oviduct, where fertilization occurs. The oviduct subsequently joins the vitelline duct, allowing each fertilized oocyte to converge with a single vitellocyte within the ootype, the specialized region of the female reproductive ducts where eggs are formed, which is surrounded by a cluster of unicellular glands (known as Mehliś gland). Eggs then pass into the uterus, which is rudimentary in mature proglottids, but expands extensively in gravid proglottids, eventually occupying most of the space while most other reproductive structures degenerate.

In this work, we describe the reproductive development of the proglottids of *H. microstoma*. We describe the development of the male and female reproductive systems, combining metabolic labeling of proliferating cells with a thymidine analog together with molecular markers of meiosis and differentiation. We show a tight coordination of development between the male and female reproductive systems, and especially of the ovary and vitelline gland within the female reproductive system, as their development occurs in rapid, almost synchronous waves of differentiation. Finally, we demonstrate the occurrence of gametogenesis and insemination in adults in an *in vitro* culture system. Our findings establish a developmental staging framework and provide an experimental system for studies of reproductive development in cestodes.

## 2. Results

Adults of the tapeworm *Hymenolepis microstoma* reach several centimeters in length and consist of hundreds of proglottids along the strobila.^23^ At the same time, the cells are very small (with nuclei typically smaller than 5 μm), making it difficult to resolve details of proliferation, differentiation, and meiotic progression by conventional histology, requiring ultrastructural analysis by electron microscopy.^16^ This challenge is compounded by the fact that major developmental transitions can occur rapidly in cestodes and may be represented by only a few consecutive proglottids along the strobila.^9^ Therefore, a comprehensive histological analysis of reproductive development throughout the entire worm is impractical.

To overcome these limitations, we chose a whole-mount approach that enabled analysis of reproductive development at each proglottid using markers of cell proliferation and differentiation. Cell proliferation was assessed by incorporation of the thymidine analogue EdU during a 2 h pulse *ex vivo* (S-phase), and by immunofluorescent detection of phospho-histone H3 (PH3), a marker of cells in mitosis and in the late stages of meiosis I and II, previously used in *Hymenolepis* spp..^26,36^ Entry of germline cells into meiosis was examined by whole-mount *in situ* hybridization (WMISH) for *hm-sycp1*, an ortholog of a conserved structural component of the synaptonemal complex found across Metazoa^37^, including flatworms^38^, which is specifically expressed during meiotic prophase I. We also analyzed the expression of other differentiation markers associated with reproductive ducts and vitellocytes. Finally, selected developmental stages were further characterized by histological analysis.

We will first separately describe the formation of the genital primordium, the development of the reproductive ducts, and the development of the male and female gonads. Finally, we will describe the temporal coordination among the different reproductive elements within the developing proglottids.

### 2.1 Initial development of the genital primordium and development of the reproductive ducts

In the earliest immature proglottids, the genital primordium is visible as a central accumulation of cells within the medullary parenchyma, which includes proliferating cells that incorporate EdU (EdU+ cells) (Figure 1A, Figure 2A). The genital primordium is surrounded by additional proliferating cells that are located adjacent to the inner longitudinal musculature (thus distributed as a ring of proliferating cells in cross sections^39^, a pattern conserved in many other cestodes^40^). Proliferating cells in the medullary parenchyma are extremely abundant in the neck region, but their abundance gradually decreases along the strobila, until no further proliferation is observed in the very last gravid proglottids.

**Figure 1.**
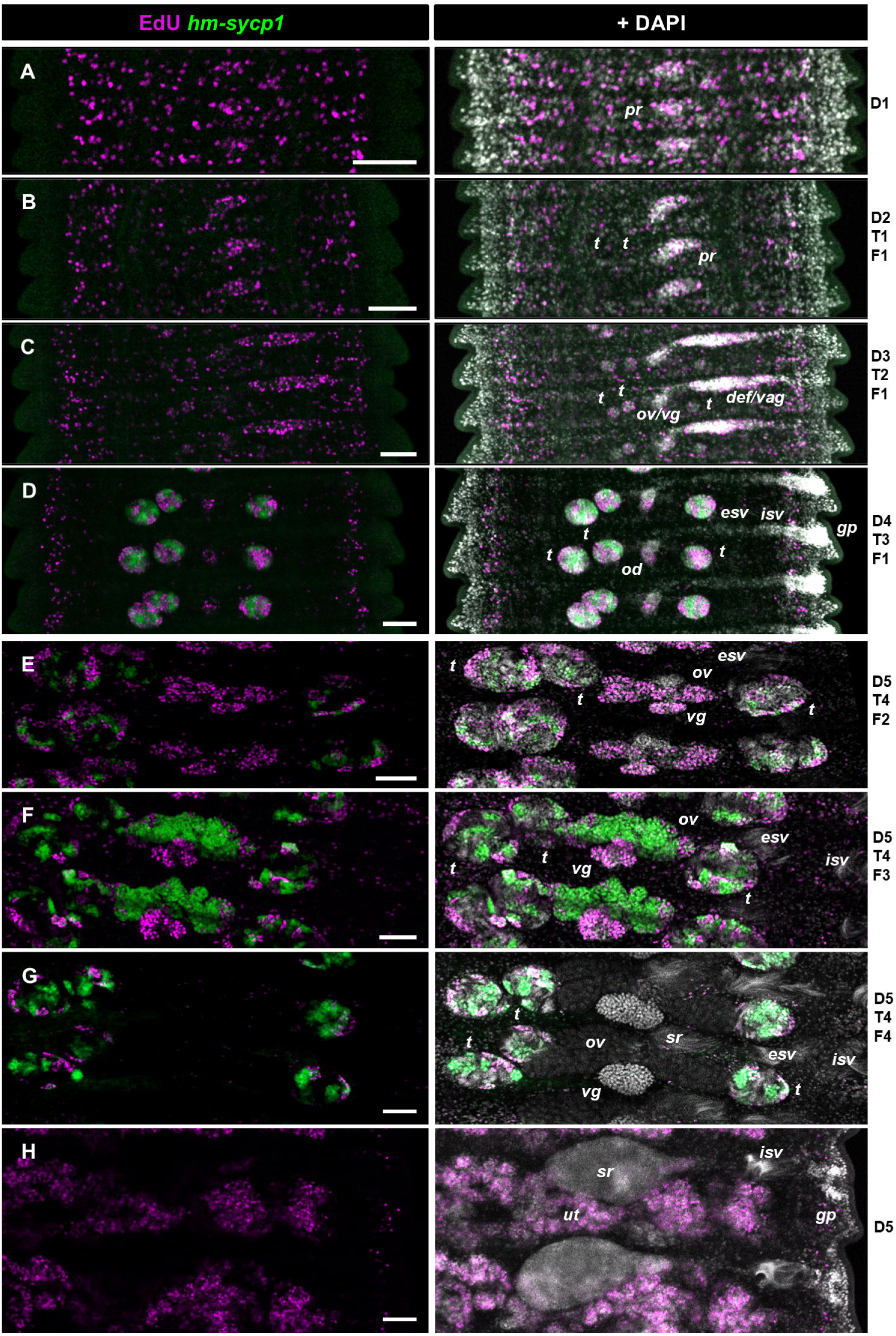
Overview of reproductive development in the strobila of *H. microstoma*. Proglottids along the strobila (**A-H**) are shown combining the detection of EdU incorporation and *hm-sycp1* expression. The stages of development (Table 1) of the reproductive ducts (D1-D5), testes (T1-T4) and ovary and vitelline gland (F1-F5) are indicated for each panel on the right. Bars: 50 μm. Abbreviations: esv, external seminal vesicle; isv, internal seminal vesicle; def/vag, primordium of the vas deferens and vagina; od, oviduct; ov, ovary; ov/vg, primordium of the ovary and vitelline gland; pr, genital primordium; sr, seminal receptacle; t, testis; ut, uterus containing embryos; vg, vitelline gland.

**Figure 2.**
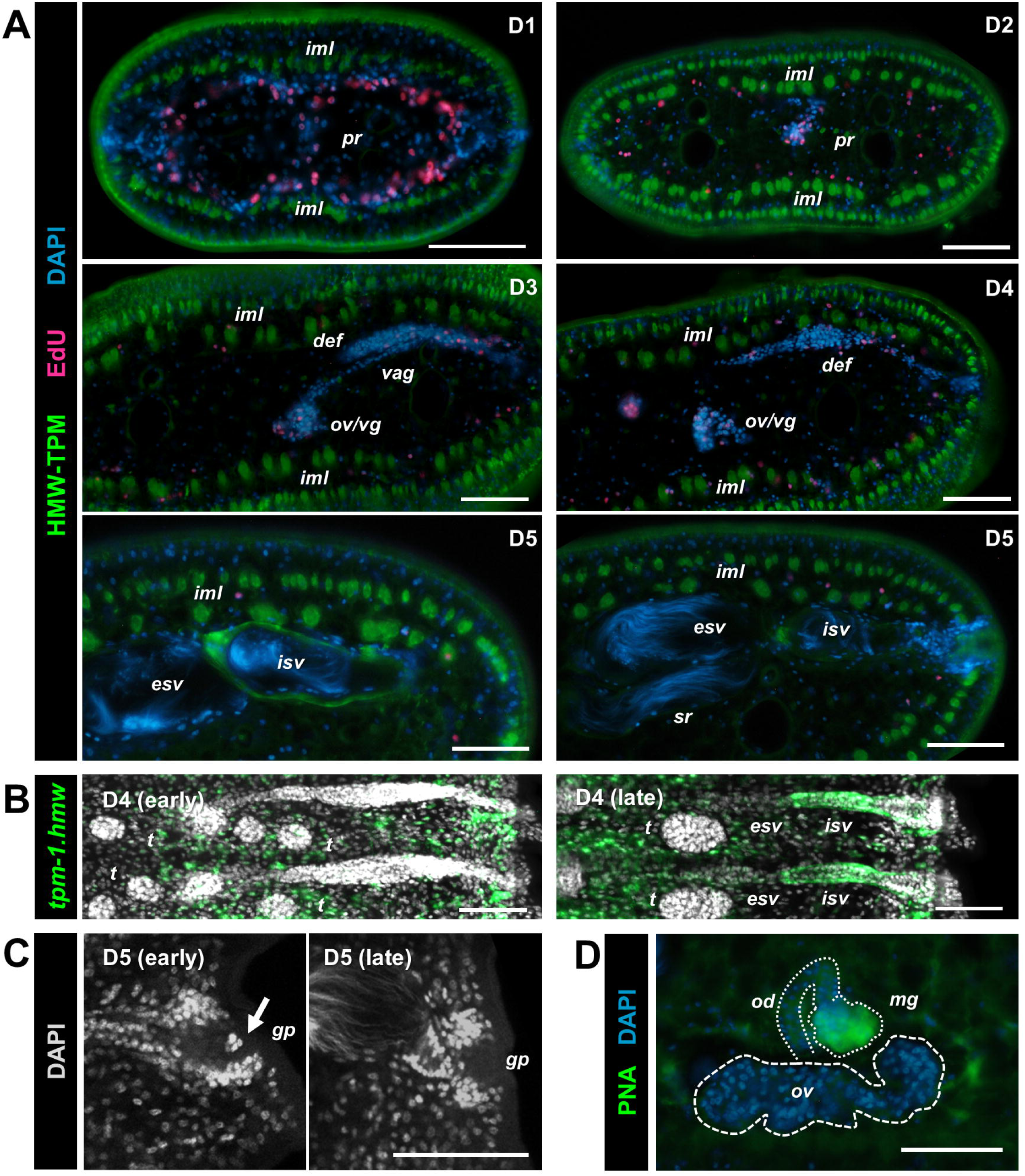
Early development of the genital primordium and development of the reproductive ducts. **A.** Successive stages of development (D1-D5, as defined in Table 1) of the genital primordium and of the reproductive ducts illustrated in histological sections stained by immunofluorescence for muscular tropomyosin isoforms (HMW-TPM) and detection of EdU incorporation. The two bottom panels in stage D5 show the ducts before (left) and after (right) insemination. **B.** Changes in the expression of the mRNA encoding the muscular tropomyosin isoform *hm-tpm-1.hmw* associated with the development of the cirrus pouch, comparing early D4 stage (incipient differentiation of internal and external seminal vesicles as dilations of a solid cord of cells) and late D4 stage (well defined internal and external seminal vesicles as hollow ducts). **C.** Disappearance of a mass of cells located at the entrance of the genital pore (arrow) at later stages of development. **D**. Labeling of the differentiated Mehliś gland by the lectin PNA. Bars: 50 μm. Abbreviations: def, vas deferens; esv, external seminal vesicle; gp, genital pore; iml, inner muscle layer; isv, internal seminal vesicle; mg, Mehliśgland; od, oviduct; ov, ovary; ov/vg, primordium of the ovary and vitelline gland; pr, genital primordium; sr, seminal receptacle; t, testis; vag, vagina;

**Table 1.** Developmental stages of the reproductive ducts: D1: central round genital primordium. D2: initial extension of a cord of proliferative cells towards the right margin, producing a comma-shaped to pistol-shaped primordium. D3: further extension of the cord of cells, producing a rifle-shaped primordium that divides into vas deferens and vagina primordia; proliferative cells restricted to the external-most region. D4: the cord of cells reaches the right margin and differentiates into the genital atrium and genital pore, without detectable cell proliferation. The vas deferens dilates to begin the formation of the external and internal seminal vesicles. Development of the oviduct and vitelloduct. D5: fully differentiated reproductive ducts, with sperm present in seminal vesicles. **<u>Developmental stages of the testes:</u>** T1: early testes primordia with few cells and sparse cell proliferation (ca. 10 μm diameter). Weak expression of *hm-sycp1* in a central cell in each testis. T2: late testes primordia, with abundant cell proliferation (ca. 20 μm diameter). Strong expression of *hm-sycp1* in a central cell in each testis. T3: testes with EdU-positive rosettes and *hm-sycp1*-positive rosettes of primary spermatocytes (ca. 30 μm diameter). T4: testes containing spermatids undergoing spermiogenesis (first detected in testes ca. 60 μm diameter). Testes continue to grow up to 100-120 μm in diameter and to produce sperm until they eventually disappear in gravid proglottids. **<u>Developmental stages of the ovary and vitelline gland:</u>** F1: ventral cell mass derived from the central genital primordium, with sparse cell proliferation and *hm-sycp1* expression, eventually divides into a recognizable ovary primordium (<70 μm in width) and vitelline gland primordium (<40 μm in width). F2: increased EdU incorporation and growth of the ovary primordium (from ca. 70 to 140 μm width) and of the vitelline gland primordium (up to 60 μm width). F3: wave of *hm-sycp1* expression in the ovary as oocytes enter meiotic prophase I (ovary ca. 140-200 μm wide). Extensive cell proliferation detectable the vitelline gland. F4: expression of *hm-sycp1* disappears from the ovary, as oocytes become arrested in diplotene I. Oocyte nuclei decondense as they grow (ovary up to 260 μm wide). Vitelline gland cell proliferation ends and vitellocyte differentiation begins (as evidenced by PNA staining). F5: oocytes are released and fertilized. The ovary and vitelline gland become depleted and disappear as the expanding uterus fills with eggs containing developing embryos.

As development proceeds, the genital primordium enlarges and becomes comma-shaped as some cells extend toward the right lateral margin of the proglottid, forming a cellular cord rich in EdU+ cells (Figures 1B, 1C, 2A). Initially, this structure consists of a single solid cord. As elongation continues, the shape first becomes similar to a pistol, and then to a rifle. At the same time, it becomes subdivided dorso-ventrally into two parallel cords that will give rise to the vas deferens (dorsally) and vagina (ventrally) (Figure 2A). During this process, proliferating cells become restricted to the most recently extended region of the cords (closer to the body wall margin) and become confined to their periphery.

At this stage, small testes primordia are already visible at the sides of the central genital primordium (Figure 1C). Developing efferent ducts extend from the testes and connect to the developing vas deferens. Continued extension of the developing vas deferens and vagina eventually brings them in contact with the body wall, where a mass of cells accumulates (Figures 1D, 2A). This structure later differentiates into the genital atrium and genital pore. Concurrently, the vas deferens develops two subtle dilatations corresponding to the future internal and external seminal vesicles. Expression of the mRNA for prohormone convertase 2 (*hm-pc2*) begins in cells surrounding the external seminal vesicle (Supplementary Figure 1A), corresponding to the developing prostatic glands.^41^

After the primordia of vas deferens and vagina reach the body wall, proliferating cells are no longer detected within them, although differentiation remains incomplete and the structures are still solid cellular cords rather than hollow ducts (Figure 1D, 2A). At later stages, muscle fibers differentiate in the cirrus sac and around the reproductive ducts, coinciding with the expression of muscle-specific tropomyosin isoforms at the mRNA and protein levels (*hm-tpm-1.hmw* and HMW-TPM, respectively)^39,42^ (Figures 2A, 2B). The proximal region of the vagina dilates and develops into the seminal receptacle. After development of the reproductive ducts is apparently complete, the genital pore remains closed by a mass of cells. This mass disappears only at later stages, after the testes have fully developed and spermatozoa have begun to accumulate within the seminal vesicles (Figure 2C).

The central region of the genital primordium gives rise to the ovary and vitelline gland in its ventral region (initially as a single primordium that eventually divides into the anterior ovary and posterior vitelline gland), and to the proximal female reproductive ducts (including the oviduct and vitelline duct). Development of the proximal female ducts occurs later than that of the vagina and vas deferens, and proliferating cells remain associated with their development after proliferation is no longer detected in the distal ducts (Figure 1D). Eventually, muscle fibers also differentiate around these ducts, and the differentiation of Mehlis’ gland becomes evident by staining with the lectin Peanut Agglutinin (PNA) (Figure 2D).

We previously reported segmental expression of the Wnt receptor *hm-fzd5/8* in the body wall during strobilation^39^. Here, we show that *hm-fzd5/8* is also dynamically expressed during the development of the reproductive ducts (Supplementary Figure 1B). Weak expression is initially detected in the developing efferent ducts and elongating genital primordium. Following the subdivision of the dorsal vas deferens and ventral vaginal primordia, expression becomes restricted to the developing internal seminal vesicle, vagina, seminal receptacle, and common genital atrium. These patterns suggest a role for Wnt signaling in the morphogenesis of the reproductive ducts.

### 2.2. Development of the testes

The testes are first distinguishable as three small clusters (two aporal and one poral) of fewer than ten cells each (Figure 1C). At these early stages, relatively little proliferation is detected by EdU incorporation. As the testes grow, however, numerous EdU+ cell patches appear (Figure 1D). Although the boundaries between individual spermatogonial rosettes are difficult to define using EdU labeling alone, PH3 immunostaining revealed isolated mitotic rosettes of two, four and eight cells (because of the much shorter duration of M phase relative to S phase, PH3 labels a much smaller proportion of the proliferative rosettes). To identify the onset of meiosis, we selected SYCP1, a conserved structural component of the synaptonemal complex that is expressed during meiotic prophase I in many animal species. The genome assembly of *H. microstoma* contains three nearly identical clustered *sycp1* genes, with 100% identity across the full coding sequence, including the region targeted by our probe. We therefore refer to their collective expression as *hm-sycp1*. The pattern observed in mature gonads (described below) supports a predominantly meiotic expression for this gene. However, we unexpectedly also detected *hm-sycp1* expression in early gonadal primordia, including the developing testes, ovary, and vitelline gland (Figure 3). This expression was observed in six different experiments, starting from different batches of adult worms.

**Figure 3.**
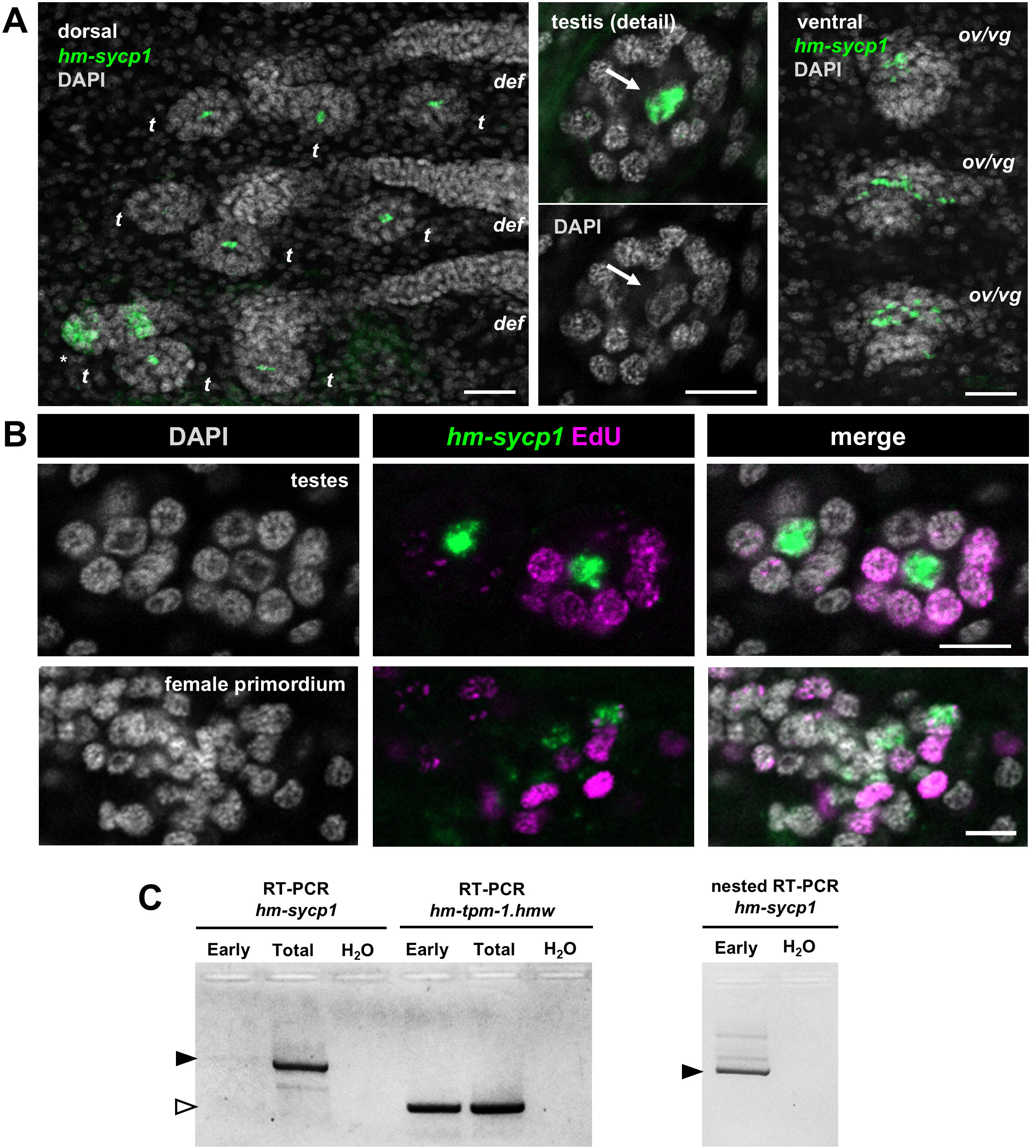
Early (pre-meiotic) expression of *hm-sycp1*. **A.** Overview of the expression of *hm-sycp1* in the primordia of testes (left and middle panels, dorsal planes) and in the primordium of the ovary and vitelline gland (right panel, ventral planes). The asterisk in the left panel indicates the first testis in the strobila in which meiotic expression of *hm-sycp1* in rosettes was detected. The arrow in the middle panel points at the single *hm-sycp1* cell in that testis primordium. **B.** Detection of EdU incorporation and *hm-sycp1* expression in early testes primordia and female primordium. **C.** RT-PCR and nested RT-PCR for *hm-sycp1* (amplification products indicated with filled arrowheads) in early pre-meiotic proglottids (“Early”) and in total strobilae (“Total”). No template control is indicated as “H_2_O”. Amplification of the muscular tropomyosin isoform *hm-tpm-1.hmw* (which is expressed in all proglottids^39^) was used as a positive control (the amplification product is indicated with an empty arrowhead). Bars: 50 μm (A, left and right panels) or 5 μm (A, middle panel, and B). Abbreviations: def, vas deferens; ov/vg, primordium of the ovary and vitelline gland; t, testis.

In the testes, the early primordia contain a single weakly positive *hm-sycp1*+ cell (Figures 3A, 3B). As the primordium enlarges, this single positive cell in each testis persists as its *hm-sycp1* signal becomes stronger, and becomes centrally located and morphologically distinct from the surrounding cells, possessing a larger polygonal nucleus with decondensed chromatin (nucleus size 6.3 ± 0.7 μm x 4.6 ± 0.5 μm, n=15) in comparison with the smaller, rounder nuclei of the cell that surround it (nucleus size 4.4 ± 0.4 μm x 3.8 ± 0.4 μm, n=45). No EdU incorporation was detected in this cell (54 analyzed testes from three worms) (Figure 3B). This *hm-sycp1*+ cell remains present when the first *hm-sycp1*+ primary spermatocyte rosettes appear (each containing 16 nuclei), which occurs when the testes reach diameters greater than approximately 30 μm (major axis 36 ± 5 μm, minor axis 27 ± 4 μm, average and standard deviations of 17 testes measured from the analysis of 5 adult worms) (Figure 3A). Beyond this stage, the fate of the central *hm-sycp1*+ cell cannot be determined because it becomes impossible to trace among the large number of spermatocyte rosettes expressing *hm-sycp1*.

Given the unexpected expression of *hm-sycp1* in early male and female gonadal primordia, we confirmed its transcription in immature proglottids by reverse transcription polymerase chain reaction (RT-PCR) analysis of anterior strobilar fragments (containing testis primordia smaller than 20 μm in diameter, and therefore lacking meiotic *hm-sycp1* expression). Conventional RT-PCR resulted in only a very weak amplification product. However, robust amplification was observed by nested RT-PCR, supporting the existence of low levels of *hm-sycp1* transcription during early gonadal development (Figure 3C). Primary spermatocytes expressing *hm-sycp1* are typically EdU-negative, as expected for cells undergoing meiotic prophase I. In some cases, however, *hm-sycp1*+ primary spermatocytes contained discrete EdU+ specks within their nuclei, possibly reflecting DNA synthesis associated with meiotic recombination (Figure 4A). Expression of *hm-sycp1* subsequently decreases and disappears during the formation of rosettes of 32 secondary spermatocytes. These give rise to rosettes of 64 rounded spermatids, whose nuclei progressively elongate during spermiogenesis. The testes continue to increase in size, eventually reaching up to approximately 100 μm in diameter. In mature testes, isolated EdU+ cells and EdU+ rosettes of 2, 4, and 8 spermatogonia are still observed, interspersed between spermatocytes and spermatids (Figure 4B, C), although more infrequently. These proliferative cells tend to be located near the periphery of the testes, whereas secondary spermatocytes, spermatids, and spermatozoa are predominantly found in the interior (Figure 4B). However, this spatial organization is less regular than that reported for other flatworms, such as planarians.^43^ Testes exhibiting *hm-sycp1* expression and EdU incorporation persist even after the uterus begins to expand, but they become progressively smaller and eventually disappear before apolysis occurs.

**Figure 4.**
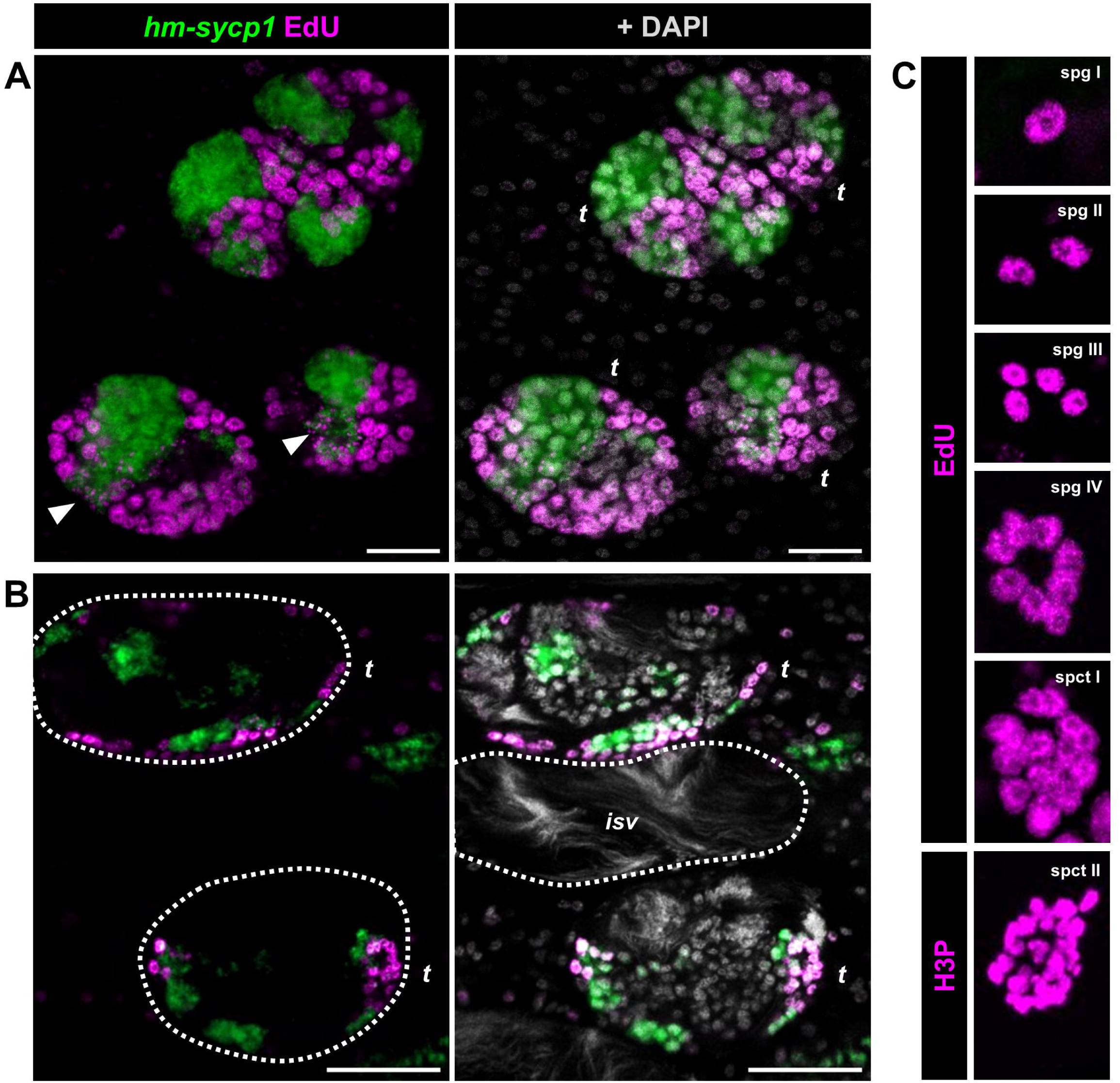
Development of the testes. **A.** Early meiotic testes containing *hm-sycp1*-expressing primary spermatocytes that are mostly EdU-negative. Some *hm-sycp1*+ primary spermatocytes contain discrete nuclear EdU+ specks (white arrowhead). **B.** Mature testes containing secondary spermatocytes and rounded and elongated spermatids (all EdU-negative) in the central testicular region. EdU+ spermatogonia and spermatocytes remain at the testis periphery. **C.** Representative stages of spermatogenesis, including EdU+ rosettes of spermatogonia I–IV, EdU+ primary spermatocytes, and H3P+ secondary spermatocytes. Bars: 20 μm. Abbreviations: t, testis; isv, internal seminal vesicle; spg, spermatogonia; spct I, primary spermatocytes; spct II, secondary spermatocytes.

### 2.3. Development of the ovary and the vitelline gland

Initially, the ovary and vitelline gland arise from a common ventral primordium, and they gradually enlarge and become morphologically distinct (Figures 1C, 5A). During this prolonged early growth phase, both organs exhibit moderate levels of EdU incorporation and occasional PH3 positive cells. In these early premeiotic stages, scattered cells expressing low levels of *hm-sycp1* are also present in both the ovary and vitelline gland. These cells appear long before the onset of meiosis, and only rarely incorporate EdU (Figures 3A, 3B, 5A, 5B).

Subsequently, both organs undergo rapid developmental changes within a relatively short interval of consecutive proglottids. In the ovary, numerous cells become EdU+, coinciding with a phase of rapid growth and likely reflecting, at least in part, widespread entry into the premeiotic S phase (Figure 5C). This is followed by a wave of *hm-sycp1* expression associated with meiotic prophase I entry, which begins when the ovary reaches a size of 140 ± 18 μm (width) x 38 ± 7 μm (length) (average and standard deviation from five worms). Expression first appears in the center of the ovary and then progressively spreads toward the periphery (Figures 5C, D and E). Cells expressing *hm-sycp1* were invariably EdU negative, and possess nuclei with condensed chromatin, consistent with meiotic prophase I. Expression of *hm-sycp1* is subsequently lost in oocytes in a wave that likewise progresses from the center toward the periphery of the ovary (Figures 5E, 5F). Following the loss of *hm-sycp1* expression, the oocytes remain arrested in meiotic prophase I, likely at the diplotene stage, and undergo dramatic growth. These oocytes have large nuclei (approximately 10 μm in diameter) containing highly decondensed chromatin (Figures 5E, 5F). The increase in oocyte size results in a concordant enlargement and weak lobulation of the ovary, reaching a size of up to approximately 260 μm (width) x 60 μm (length). The approximate total number of oocytes produced in each proglottid was estimated to be between 400 and 500, based on manual counting of oocytes in confocal stacks of two mature ovaries.

**Figure 5.**
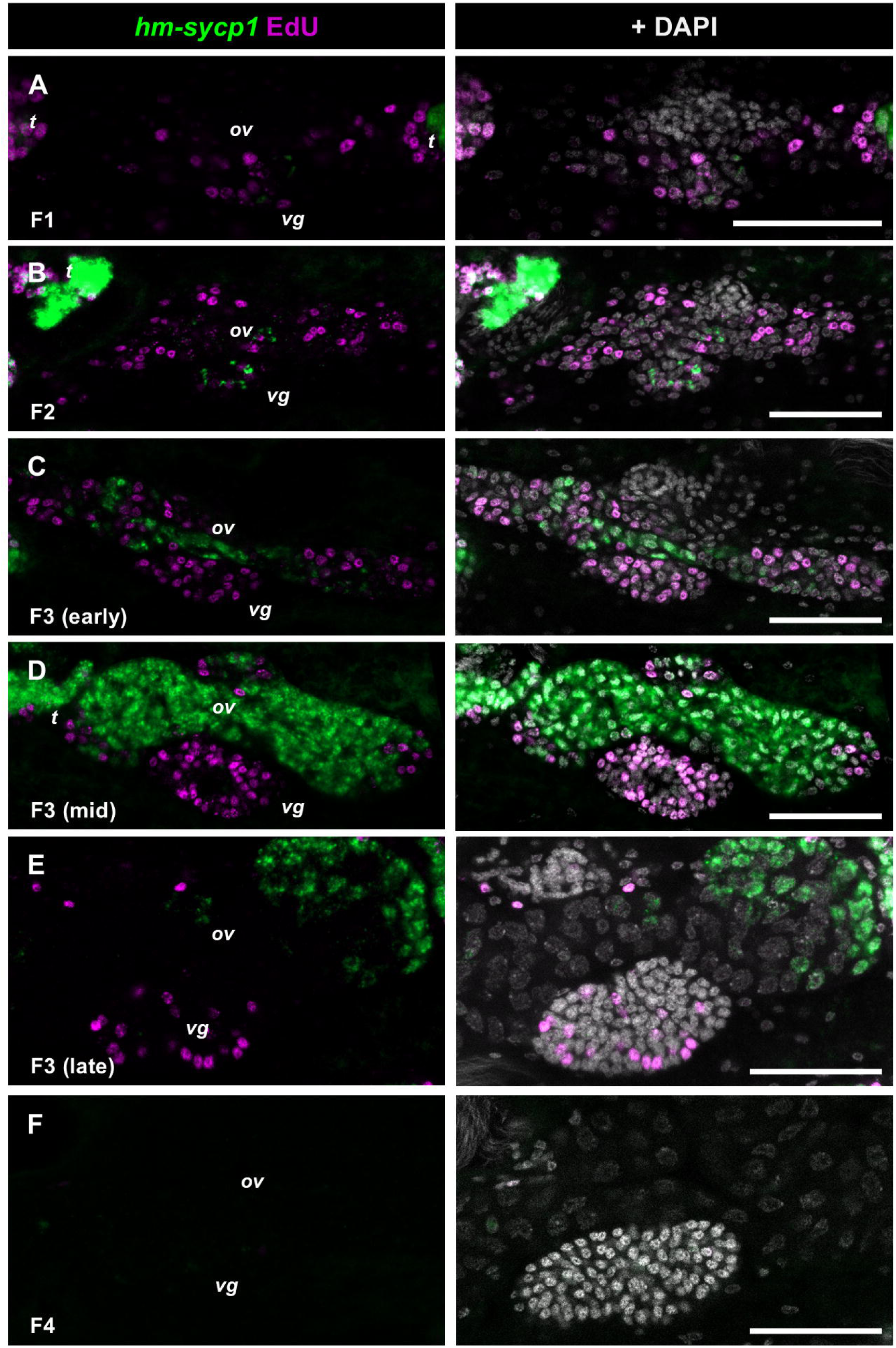
Development of the ovary and vitelline gland. Successive stages (**A-F**) are shown combining the detection of EdU incorporation and *hm-sycp1* expression. The corresponding developmental stages (F1-F4, as defined in Table 1, with the additional differentiation of early, mid and late F3 stage) are indicated on the bottom left of each panel. Bars: 50 μm. Abbreviations: ov, ovary; t, testis; vg, vitelline gland.

The development of the vitelline gland occurs in parallel with meiotic entry in the ovary. During the period when ovarian cells express *hm-sycp1*, the vitelline gland exhibits extensive proliferation, with most cells incorporating EdU after 2 h of labelling (61 ± 11% of all cells in the gland, average and standard deviation of six glands, with 29-80 cells analyzed per gland) (Figures 5C, 5D). Strikingly, termination of proliferation in the vitelline gland coincides with the loss of *hm-sycp1* expression in the ovary. Both transitions occur within fewer than 20 consecutive proglottids, from the point of maximum *hm-sycp1* expression in the ovary and EdU incorporation in the vitelline gland, until *hm-sycp1* expression in the ovary and EdU incorporation in the vitelline gland become undetectable (Figures 5D, 5E, 5F). This indicates a tight temporal coordination between ovarian and vitelline gland development. Following the end of proliferation, vitelline cells continue their differentiation. Immediately after their proliferation ceases, vitelline cells are negative for PNA lectin staining, but subsequently become strongly PNA-positive (Figure 6A), consistent with the previous description of differentiated vitellocytes in *Hymenolepis diminuta*.^26^ A similar pattern is observed for the expression of *hm-zf621400*, a gene previously identified as a marker of the vitelline gland.^27^ Expression of *hm-zf621400* is absent from vitelline glands containing proliferating cells, and weak expression begins after proliferation has ceased. Strong expression is observed in differentiated vitellocytes that are released to accompany fertilized oocytes (Figure 6B). Unexpectedly, weak *hm-zf621400* expression is also detected in the mature ovary, particularly in oocytes adjacent to the vitelline gland. In addition, expression is present in parenchymal cells, as previously reported by Olson et al.^27^ (not shown).

**Figure 6.**
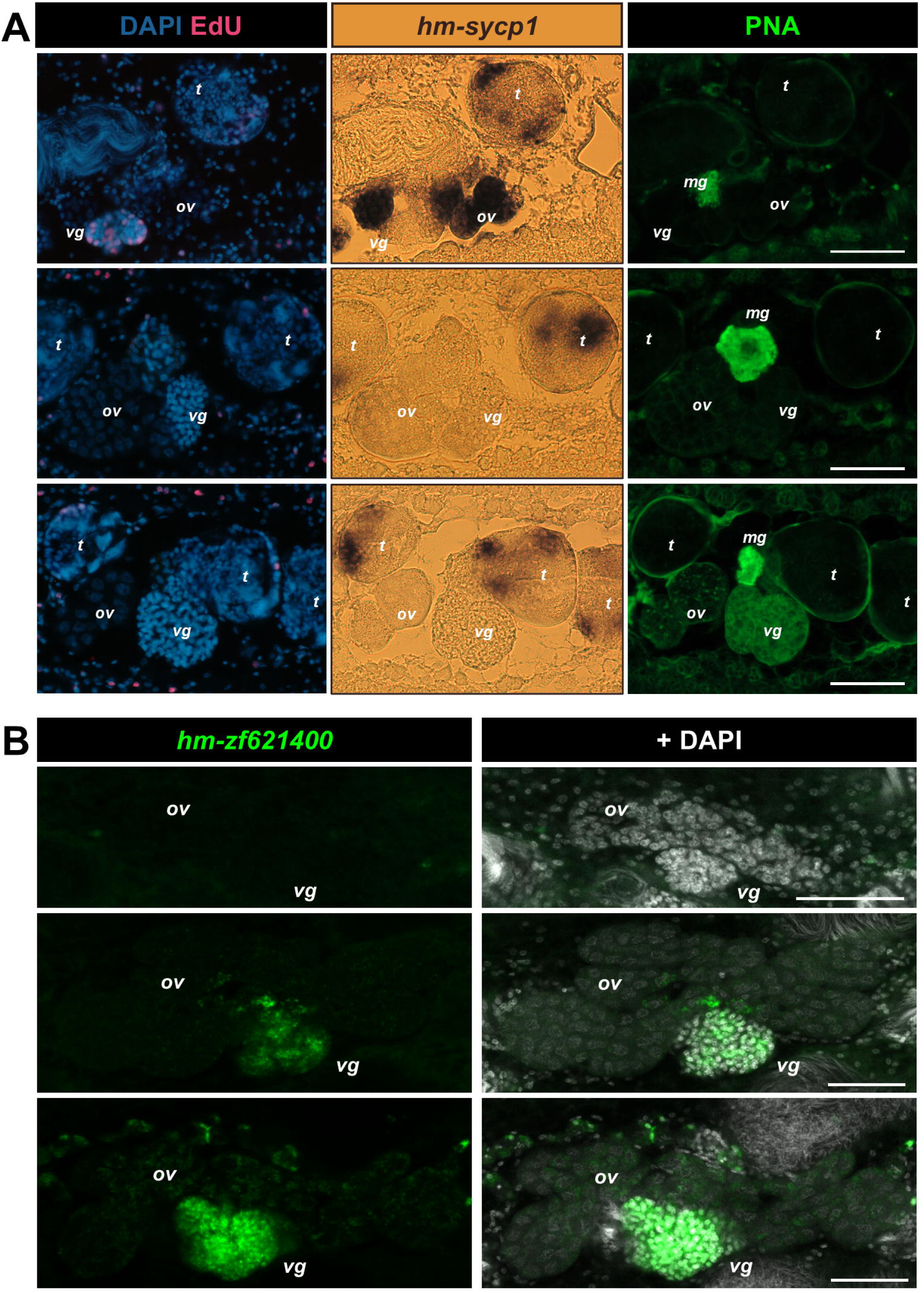
Differentiation of the vitelline gland. **A.** Histological sections of proglottids stained for the detection of EdU incorporation, *hm-sycp1* expression, and PNA labeling. Top row, proglottid with ongoing meiotic expression of *hm-sycp1* in the ovary and proliferation in the vitelline gland. Middle row, proglottid found immediately after meiotic expression of *hm-sycp1* in the ovary disappears and proliferation in the vitelline gland ends. Bottom row, later proglottid with ovary containing mature oocytes and differentiating vitelline gland (labeled with PNA). **B.** Expression of *hm-zf621400* during the development of the female reproductive system. Top row, F2 stage. Middle row, early F4 stage. Bottom row, late F4 stage. Bars: 50 μm. Abbreviations: mg, Mehliśgland; ov, ovary; t, testis; vg, vitelline gland.

### 2.4. Coordination of male and female reproductive development

To compare developmental progression among each component of the reproductive system, we established independent staging schemes combining morphological and molecular markers for the reproductive ducts (D1–D5), testes (T1–T4), and ovary and vitelline gland (F1–F5) (Table 1). We then scored each proglottid according to all three schemes and analyzed the combinations of stages observed along the strobila.

Detailed analyses of four adult worms obtained from three different experiments evidenced the differences in the dynamics of male and female reproductive development (Figure 7). Most proglottids in the strobila corresponded to stages characterized by formation of the genital primordium and reproductive ducts, together with testicular growth preceding spermiogenesis, suggesting that these phases occupy a relatively long developmental period. In contrast, female reproductive development was concentrated within a much shorter region of the strobila, although the absolute number and proportion of proglottids at each stage varied among individuals. Notably, the onset of rapid ovarian and vitelline differentiation coincided with the appearance of testes undergoing spermiogenesis and with the accumulation of spermatozoa in the seminal vesicles.

**Figure 7.**
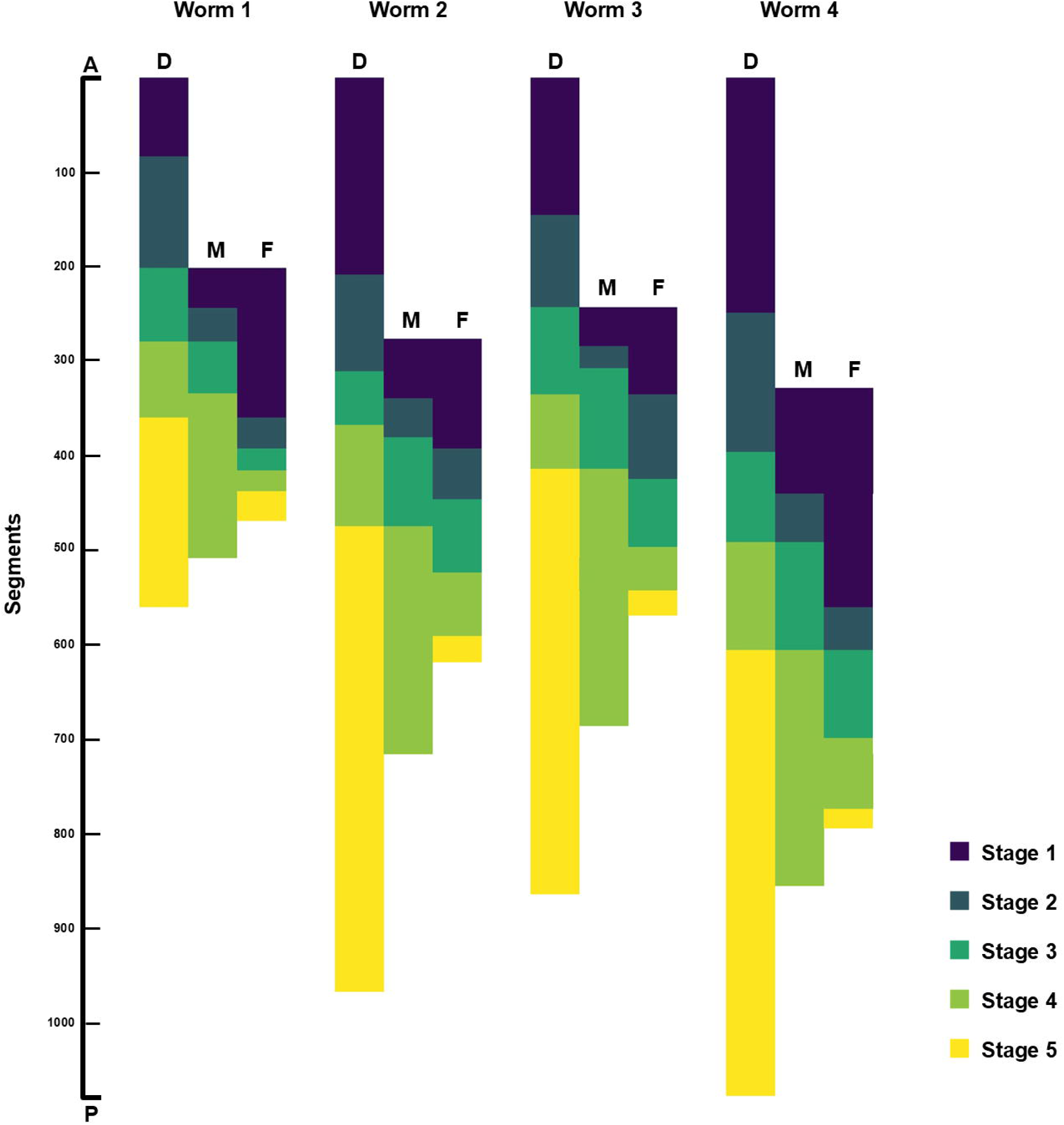
Coordination of the development of the ducts (D), male reproductive system (M), and female reproductive system (F). Distribution of developmental stages (Table 1) for each system along the strobila (proglottids) of four individual adult worms (worms 1–4) stained for the detection of EdU incorporation and *hm-sycp1* expression. Proglottids are arranged along an anterior–posterior gradient, from the anterior region (A), containing the least developed reproductive systems, to the posterior region (P), containing the most mature proglottids.

The female seminal receptacle gradually accumulated spermatozoa during subsequent stages, suggesting repeated or continuous insemination events (Figures 1G, 1H). Determining the precise onset of insemination from whole-mount preparations proved difficult because the external and internal seminal vesicles rapidly become filled with spermatozoa, obscuring the adjacent vagina and seminal receptacle. To resolve this issue, we analyzed serial histological sections of strobilar fragments of worms processed either for *hm-sycp1* WMISH and EdU labeling, or for immunofluorescence for muscular tropomyosin isoforms (HMW-TPM)^42^ and EdU labeling. Similar results were observed in a worm stained as a whole-mount for actin with phalloidin (Figure 8A), which better revealed the position of each reproductive duct as it stained the thin muscle fibers that surround them (although it is unfortunately incompatible with our EdU detection and WMISH protocols).

**Figure 8.**
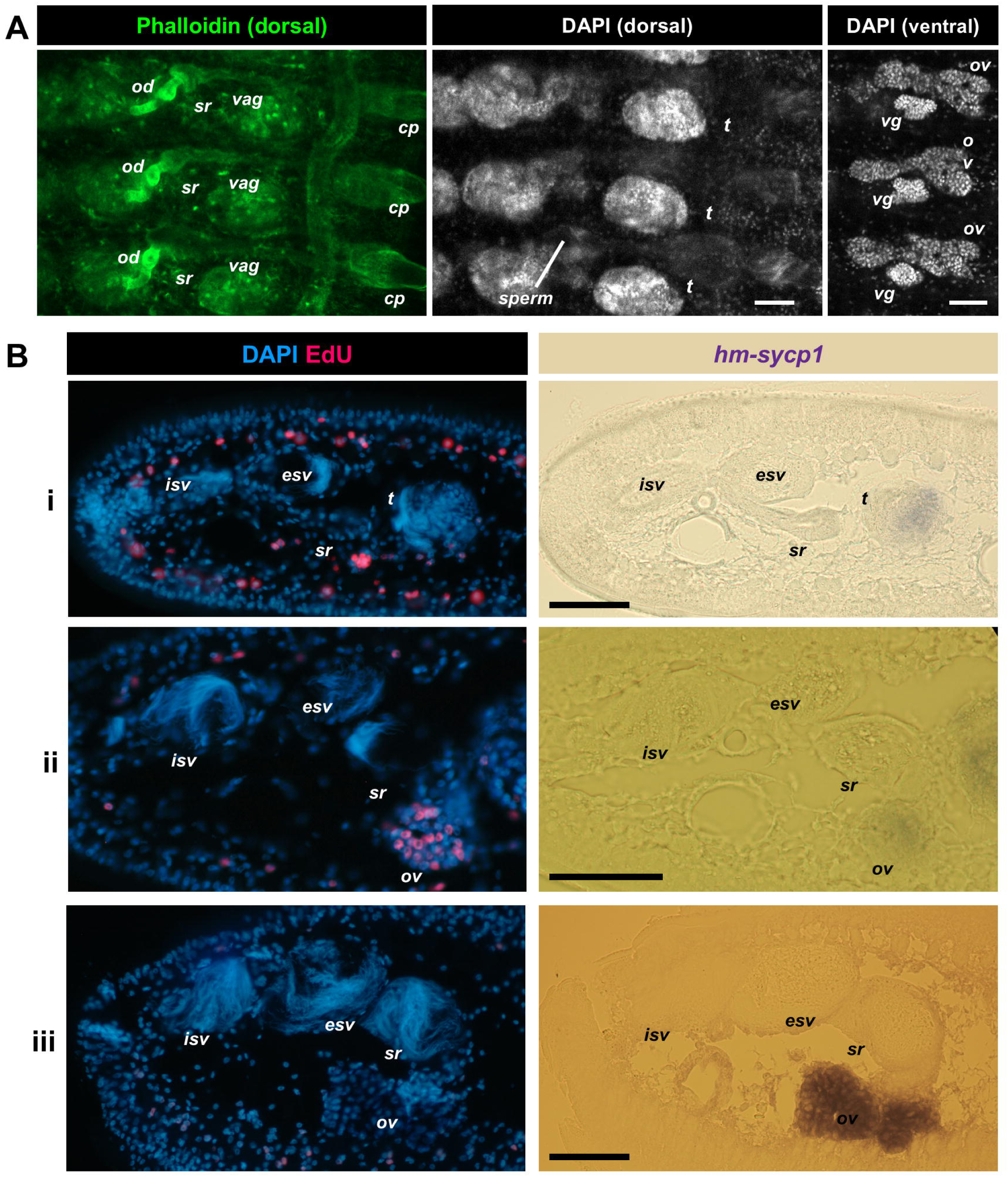
Insemination. Identification of the earliest proglottids containing spermatozoa in the seminal receptacle in **(A)** a complete strobila stained with phalloidin and DAPI, and **(B)** serial sections of a strobila stained for the detection of EdU incorporation and *hm-sycp1* expression. In **A**, the left and middle panels show the dorsal region, and the right panel shows the corresponding ventral region of the same proglottids. In **B**, the rows show **i)** a proglottid immediately before insemination, **ii)** an example of one of the first proglottids after insemination (containing a small amount of sperm in the seminal receptacle), and **iii)** a later proglottid with a seminal receptacle filled with spermatozoa. The distance between ii and iii is approximately four proglottids. Bars: 50 μm. Abbreviations: cp, cirrus pouch; esv, external seminal vesicle; isv, internal seminal vesicle; od, oviduct; ov, ovary; sr, seminal receptacle; t, testis; vag, vagina; vg, vitelline gland.

These analyses showed that insemination began shortly after spermatozoa first appeared within the male reproductive ducts. In all three individuals examined in detail, the earliest spermatozoa detected in the seminal receptacle were observed to approximately coincide with the entry of oocytes into prophase I (as determined by *hm-sycp1* expression, loss of EdU incorporation, and ovary size) (Figure 8B). This finding indicates a close temporal coordination between male reproductive maturation, insemination, and female meiotic development.

After both female and male reproductive systems are mature, there is a sharp transition in the strobila as oocytes are quickly fertilized and incorporated into eggs. Thus, the ovary and vitelline gland are quickly spent and disappear, and proglottids containing mature testes and a mature ovary comprise a very small proportion of the strobila (fewer than 10% of all proglottids, n=4 worms). The testes continue to be present and to produce sperm in early gravid proglottids (as evidenced by continued expression of *hm-sycp1* and EdU incorporation) until they also disappear in late gravid proglottids. The male distal ducts (vas deferens with seminal vesicles) and the female distal ducts (vagina and seminal receptacle) are not eliminated and remain filled with spermatozoa until the last gravid proglottids.

### 2.5. Gametogenesis *in vitro*

Finally, we investigated whether gametogenesis could proceed in adult *H. microstoma* maintained *in vitro*. Using a culture system based on methods described by Seidel^44^, we performed EdU pulse-chase experiments to follow the differentiation of male and female germ cells (Figure 9). Two independent experiments yielded similar results.

**Figure 9.**
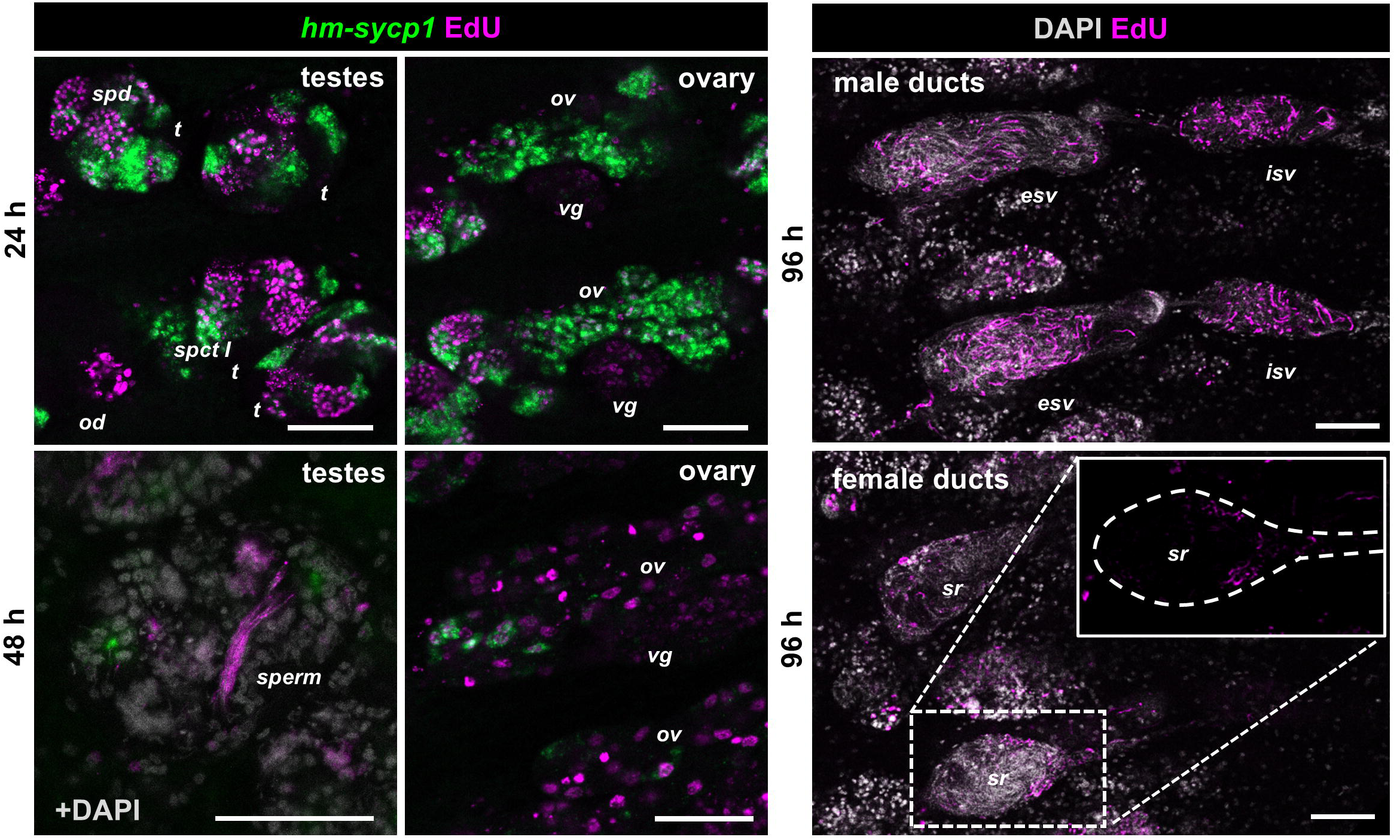
Detection of gametogenesis by EdU pulse-chase *in vitro*. Worms were incubated for 2 h in media with EdU, chased in EdU-free media for the intervals of time indicated next to each panel, and stained to detect EdU and *hm-sycp1* expression. Bars: 50 μm. Abbreviations: esv, external seminal vesicle; isv, internal seminal vesicle; ov, ovary; spct I, primary spermatocytes; spd, spermatids; sr, seminal receptacle; t, testis; vg, vitelline gland.

In the testes, a 2 h EdU pulse labeled spermatogonia and primary spermatocytes prior to meiotic entry, as described above. After a 24 h chase in EdU-free medium, labeled cells were detected among secondary spermatocytes and round spermatids. Following 48 h of chase, EdU labeling was observed in elongating spermatids within the testes, whereas after 96 h labeled spermatozoa were present in the external and internal seminal vesicles. In addition, labeled spermatozoa were detected in the seminal receptacle of some proglottids, demonstrating insemination with spermatozoa that differentiated during *in vitro* culture.

In the ovary, a 2 h EdU pulse labeled oogonia and *hm-sycp1* negative oocytes prior to meiotic entry. After a 24 h chase, EdU and *hm-sycp1* signals co-localized within the same oocytes, indicating entry into meiotic prophase I during culture. After 48 h, large EdU-labeled oocytes lacking *hm-sycp1* expression were observed, corresponding to oocytes arrested in meiotic prophase I after *hm-sycp1* is turned off, as described above.

Together, these results demonstrate that both male and female gametogenesis continued in adult worms maintained *in vitro*. Furthermore, the appearance of newly generated (EdU+) spermatozoa within the seminal receptacle indicates that insemination can also occur under these culture conditions.

## 3. Discussion

### 3.1. Variation in strobilar development

The total number of proglottids, as well as the number of proglottids assigned to each developmental stage, varied substantially among individuals despite the use of a single parasite strain maintained in an inbred mouse line under standardized laboratory conditions. The extent of variation present in natural populations is likely to be substantially greater. Although the strobila provides a remarkably complete developmental series, it is not clear whether the number of proglottids occupying a given stage reflects the duration of that stage, as this relationship depends on whether proglottids are generated at a constant rate in the neck region, and if the rate of development of the reproductive system between proglottids is also constant. Direct measurements of the rates of proglottid production are not available for *H. microstoma*, but we reanalyzed published data describing the number of proglottids present in worms at different times following infection for the related species *Hymenolepis diminuta*^45^ (Supplementary Figure 2). Remarkably, the relationship between proglottid number and time was highly linear once strobilation began, suggesting that successive proglottids are generated at a nearly constant rate of approximately 8 min per proglottid.

Although the rate of proglottid generation has not been measured in *H. microstoma*, the developmental timing described by de Rycke and Grembergen^35^ is consistent with our observations. The first mature proglottids appeared approximately 9 days post-infection, whereas only about 2 additional days elapsed before the ovary and vitelline gland disappeared, supporting the conclusion that female reproductive maturation occurs over a short interval. Gravid proglottids containing infective eggs were present 15 days post-infection.

Combining our data with the developmental sequence reported by de Rycke and van Grembergen^35^ allows a rough estimate of the rates of proglottid generation in *H. microstoma*. Assuming that developmental rates are similar during the initial phase of strobilar growth (described by de Rycke and Grembergen^35^) and in the patent infections examined here, the observation that approximately 4 days elapse between the appearance of the first proglottids and of the first clearly differentiated testes^35^ can be compared with our finding that 334 ± 87 proglottids separate these two developmental landmarks. This comparison yields an estimated time for the generation of each proglottid of approximately 17 min. Comparing the number of proglottids found between two other developmental landmarks (264 ± 58 proglottids from the appearance of the first clearly differentiated testes until the first embryos are generated; these stages were separated by 4 days in the work of de Rycke and Grembergen^35^), a similar estimate of 22 minutes was obtained.

### 3.2. Early reproductive development

Early reproductive development in *H. microstoma* begins with the formation of a central genital primordium. The overall pattern is similar to that described in other cestodes, including the closely related *Hymenolepis diminuta*^11^ as well as more distantly related species.^5,9,10^ From this genital primordium, a cellular cord giving rise to the reproductive ducts extends toward the region of the future genital pore.

Our observations suggest that the cells of the ovary and vitelline gland originate from the central region of the genital primordium. In contrast, the origin of the testes remains unclear. Testes may derive from cells of the genital primordium itself, as proposed by Douglas^9^ for *Cylindrotaenia diana* (syn. *Baerietta diana*), or alternatively from surrounding parenchymal cells. Distinguishing between these possibilities is difficult in the absence of markers that can identify germline cells prior to gonadal differentiation.

The early expression of *hm-sycp1* in gonadal primordia, long before the onset of meiosis, was unexpected. Although SYCP1 is best known as a component of the synaptonemal complex during meiotic prophase I, early expression of synaptonemal complex genes has also been reported in vertebrate primordial germ cells (PGCs).^46–48^ Could the early *hm-sycp1*+ cells observed in developing testes and ovaries represent primordial germ cells or early germline stem cells (GSCs)? The solitary *hm-sycp1*+ cell present in each early testis primordium occupies a central position and displays a morphology corresponding with the germinative cells described by Sulgostowska in early testes of *H. diminuta*.^11^ These germinative cells were hypothesized to represent the germ line, in contrast with other undifferentiated proliferating cells found in the testes, named as “germinative-somatic cells” in that study. However, if this cell represents a PGC or GSC, it does not appear to proliferate during the period of rapid testicular growth (as it did not incorporate EdU), and would therefore have to be a quiescent reserve population. A similar interpretation could apply to the ovary, where scattered *hm-sycp1*+ cells are present long before meiotic entry, which only rarely incorporate EdU. Interestingly, low-levels of *hm-sycp1* expression are also detected in the early vitelline gland primordium. Developmental and evolutionary evidence suggests a common origin between oocytes and vitellocytes, deriving from the ancestral flatworm female germ line, and vitellocyte progenitors display germline-associated gene expression profiles in planarians.^49^ Conversely, we observed weak expression of the vitelline marker *zf621400* in a subset of oocytes. It is possible that these shared expression patterns reveal ancestral similarities in their gene regulatory networks.

### 3.3 Coordination of male and female reproductive development

Our analyses revealed the coordination between male and female reproductive development. Insemination occurs shortly after spermatozoa first appear in the male reproductive ducts and coincides with the rapid differentiation of the female reproductive system. The coupling of extensive cell proliferation in the vitelline gland with the wave of ovarian meiotic differentiation suggest the existence of regulatory mechanisms acting at the level of the entire female reproductive system. It is conceivable that these organs could communicate directly or respond to a common developmental signal.

One possibility is that insemination itself could contribute to the regulation of female reproductive development. In many dioecious animals and cross-fertilizing hermaphrodites, seminal fluids have been shown to contain signaling molecules capable of influencing the physiology and development of the female reproductive system.^50–54^ Whether similar mechanisms operate in self-fertilizing hermaphrodites remains largely unexplored. In *H. microstoma*, insemination occurs at approximately the stage when *hm-sycp1* expression first appears in the ovary, raising the possibility that signals from spermatozoa or seminal fluid components could contribute to the regulation of development of the female reproductive system. Our previous work showed that the prostatic glands of *H. microstoma* express the prohormone convertase *hm-pc2*^41^, suggesting they may secrete neuropeptides or peptide hormones that could be released into the seminal fluid. Ultrastructural observations in *Cylindrotaenia hickmani*^55^ demonstrated that secretions from the prostatic glands became associated with the surface of spermatozoa, potentially providing a mechanism for the delivery of signaling molecules during insemination.

Interestingly, spermatozoa can be found within the reproductive ducts long after the ovary and vitelline gland have regressed. In the case of spermatozoa stored within the seminal vesicles, their persistence could simply reflect a role in inseminating non-gravid proglottids. However, the continued presence of spermatozoa within the seminal receptacle is more difficult to explain, but could be compatible with an ongoing role in the regulation of reproductive development.

### 3.4. Gametogenesis *in vitro*

Robust *in vitro* culture systems have been developed for *Hymenolepis diminuta*, supporting both larva-to-adult development and regeneration of the strobila following amputation.^7,56^ In contrast, although an *in vitro* culture system capable of supporting development of *H. microstoma* from larva to adult was reported several decades ago,^44^ these results have not been successfully replicated in more recent attempts.^25^ Here, rather than attempting the complete transition from larva to adult, we focused on the maintenance of fully developed adult worms to determine whether reproductive development and gametogenesis could continue under *in vitro* conditions.

Our pulse-chase experiments demonstrate that reproductive development remains active for at least 96 h in culture. Male germ cells progressed through spermatogenesis, newly generated spermatozoa participated in insemination, and female germ cells entered and progressed through the early stages of meiotic prophase I. The rate of spermatogenesis was slower than for *H. diminuta in vivo*^33^, suggesting that our culture conditions do not fully recapitulate the physiological environment of the host. In addition, a contribution of EdU toxicity to the observed developmental rates is possible. Although further optimization will likely improve the system, these results establish adult *in vitro* culture as a useful platform for future functional studies of gametogenesis and reproductive development in cestodes.

## 4. Experimental Procedures

### 4.1. Laboratory maintenance of *H. microstoma*

*Hymenolepis microstoma* (Nottingham strain) was maintained using *Mus musculus* (C57BL/6) as the definitive host and *Tribolium confusum* as the intermediate host, as previosuly described.^23^ Maintenance of the parasite life cycle was conducted in collaboration with the Laboratorio de Experimentación Animal, Facultad de Química, Universidad de la República, Uruguay, under protocol No. 10190000025215 (“Mantenimiento del ciclo vital completo del cestodo *H. microstoma* utilizando sus hospedadores naturales *Mus musculus* (ratón) y *Tribolium confusum* (escarabajo de la harina)”), approved by the Comisión Honoraria de Experimentación Animal, Uruguay.

Adult worms were recovered from infected mice 2–3 months post-infection, washed in phosphate-buffered saline (PBS), and either fixed overnight at 8 °C with 4% paraformaldehyde prepared in PBS for most subsequent analyses, or maintained for *ex vivo* and *in vitro* culture experiments.

### 4.2. Phalloidin staining

Fixed worms were extensively washed with PBS plus 0.3% Triton X-100 (PBS-Tx), blocked with 1% bovine serum albumin (BSA) in PBS-Tx for at least one hour at room temperature, and incubated overnight at 4 °C with phalloidin-iFluor 488 (Abcam, ab176753, U.S.A., 1:1000 dilution) and 4’,6-diamidino-2-phenylindole (DAPI, Merck, 1 μg/ml). After extensive washes with PBS-Tx, specimens were sectioned into fragments of approximately 1 cm in length and mounted with ProLong Glass mounting media (Thermo Fisher Scientific, P36982)

### 4.3. Whole-mount *in situ* hybridization (WMISH)

Partial fragments of the coding sequence of each gene were obtained by RT-PCR amplification of *H. microstoma* adult cDNA and cloning into the TA Cloning™ Kit, Dual Promoter, with pCR™ (Thermo Fisher Scientific, K207040). The primers used for *hm-sycp1* (corresponding to three genes with identical coding sequences in the *H. microstoma* genome assembly, Wormbase Parasite codes HmN_003000430, HmN_003000500, HmN_003000540)^57^ were 5’-ACATGGATCAGGATGAGTTCACAA-3’ (forward primer) and 5’-TCCACAATTGGGACATTCAAAAGG-3’ (reverse primer). The probes used for *hm-zf621400* (HmN_000621400)^27^, *hm-tpm1.hmw* (HmN_000188900)^36^ and *hm-fzd5/8* (HmN_000386300)^39^ were previously described. RNA probes labeled with digoxigenin-UTP (Merck, DIGUTP-RO) were generated by *in vitro* transcription with T7 or SP6 polymerases of cloned fragments of the coding sequence of each gene. Fluorescent WMISH and colorimetric WMISH were performed with these probes as previously described.^6^ When WMISH was combined with EdU labelling or WMIHF, these protocols were performed after WMISH was complete. Fluorescently labeled whole-mount samples were mounted as fragments of approximately 1 cm in length with ProLong Glass mounting media (Thermo Fisher Scientific, P36982) or with 80% glycerol, 50 mM Tris.HCl, pH 8.0.

### 4.4. *Ex vivo* labelling with 5-Ethynyl-2’-deoxyuridine (EdU)

Adult worms were incubated for 30 min in RPMI 1640 media, HEPES modified (Merck R4130) supplemented with 10% fetal bovine serum (Capricorn FBS-11A) at 37 °C for acclimation. EdU (Click-iT Plus EdU Alexa Fluor 555 Imaging Kit, Thermo Fisher Scientific, C10338) was then added to a final concentration of 10 μM, and worms were incubated for 2 h at 37 °C. After incubation, worms were fixed as described above, and EdU detection was carried out using the Click-iT Plus EdU Alexa Fluor 555 Imaging Kit (Thermo Fisher Scientific, C10338), or Click-iT Plus EdU Alexa Fluor 488 Imaging Kit (Thermo Fisher Scientific, C10337) following the instructions of the manufacturer. Samples were co-stained with DAPI (1 μg/mL) as a nuclear marker.

### 4.5. Whole-Mount Immunofluorescence

Prior to immunofluorescence, samples were subjected to heat-induced epitope retrieval (HIER) in a 10 mM citrate buffer (pH 6.0) at 99 °C for 20 min. Whole-mount immunofluorescence was then performed on fragments comprising approximately four proglottids from different regions of the adult strobila that had previously been processed for *in situ* hybridization.

Immunostaining was modified from Koziol et al., 2013^58^, in which all incubations with antibodies were performed for 3 days in a blocking solution with 0.02% sodium azide. Samples were incubated with a rabbit polyclonal anti-phospho-Histone H3 (Ser10) antibody (Cell Signalling Technology #9701; 1:200 dilution) for 3 days at 4 °C and were subsequently incubated with an Alexa Fluor 546-conjugated anti-rabbit IgG secondary antibody (Invitrogen, A11010; 1:500 dilution).

### 4.6. Histological sections

After fixation, adult worms (with and without EdU labeling) were cut into fragments of approximately 1 cm of length, and each fragment was dehydrated, included into Paraplast (Oxford Labware), and cut into histological sections of 10 to 20 μm in thickness. The same protocol was performed with adult worms following colorimetric WMISH staining. These sections were deparaffinized, and used for diverse staining procedures, including detection of EdU incorporation (Click-iT Plus EdU Alexa Fluor 555 Imaging Kit, Thermo Fisher Scientific C10338, as instructed by the manufacturer), immunofluorescence with a polyclonal antibody generated against muscular tropomyosin isoforms as previously described^42^, and staining with Peanut Agglutinin (PNA) lectin (Lectin Kit I, Fluorescein, Vector Laboratories, FLK-2100). PNA staining was performed by blocking the deparaffinized sections with PBS plus 0.1% Triton X-100 and 1% bovine serum albumin for one hour at room temperature, followed by overnight incubation at 4 °C in the same buffer with 10 μg/mL PNA-Fluorescein and 1 μg/ml DAPI. Samples were washed extensively with PBS plus 0.1% Triton X-100 and mounted with Fluoroshield (Merck, F6182).

### 4.7. Microscopy

Whole-mount samples were imaged by confocal microscopy (Zeiss LSM 800CyAn, Advanced Bioimaging Unit of the Institute Pasteur of Montevideo) and histological sections with a Nikon Microphot-FXA epifluorescence microscope.

### 4.8. RNA extraction and RT-PCR

The anterior strobila of each of seven live adult worms was cut into pieces of alternating sizes: pieces of only approximately 5 to 10 proglottids, which were fixed and stained as wholemounts with DAPI, alternating with larger pieces of approximately 20 to 30 proglottids, which were stored in Trizol (Invitrogen, 15596026) at -20 °C until further use. The relative position of each piece was recorded, and by observing the developmental stage of the DAPI stained fragments, we were able to select the fragments stored in Trizol corresponding to the earliest developmental stages (lacking testes or with testes primordia with a diameter of 20 μm or smaller). The selected fragments were pooled, and RNA was extracted with Direct-zol RNA Miniprep (Zymo Research). Another RNA extraction was prepared from whole adult worms as a control. RNA (500 ng) was reverse transcribed with SuperScript II reverse transcriptase (Thermo, 18064014) using oligo dT as primer. The obtained cDNA was used for RT-PCR with Mango Taq polymerase (Bioline, BIO-21082) using primers for *hm-sycp* (5’-ACATGGATCAGGATGAGTTCACAA-3’ and 5’- TCCACAATTGGGACATTCAAAAGG-3’) and for *hm-tpm-1.hmw* as a positive control (5’-AGTCGTAAGGCCCTTGAGACCAG-3’ and 5’-GAAGTGAGTTCCTCGAAAGTGGTG-3’)^36^. After an initial denaturation of the template at 94 °C for 3 min, the cycling program consisted in denaturing at 94 °C for 15 s, annealing at 55 °C for 15 s, and extension at 72 °C for 90 s, for 35 cycles. For nested RT-PCR, 0.5 μl of the first PCR reaction, or from an identical control from a no-template control PCR reaction, were used as templates for a second PCR reaction using nested primers for *hm-sycp1* (5’-ACAAATCGGAGGCCTGAATGAG-3’ and 5’-GCACACGAATTTGCGGAGTTC-3’) with identical cycling conditions except that extension was performed for 30 s. All primer pairs spanned several exons, allowing to distinguish cDNA from gDNA amplification.

### 4.9. *In vitro* culture of adult worms

Adult worms were maintained in a biphasic culture system consisting of RPMI 1640 medium (HEPES modified) supplemented with 30% fetal bovine serum (Capricorn FBS-11A), 1% antibiotic-antimycotic solution (containing Penicillin, Streptomycin and Amphotericin B, Thermo Fisher, 15240062), 0.15 μM hemin (Merck, H9039), and 0.1% sodium choleate (Merck, S9875), with a solid base of tryptone agar with 5% ovine blood (Biokey, Uruguay). Cultures were set up at 37 °C and 5% CO_2_ in upright T-25 culture flasks (two adult worms per flask), in which the solid base was laid on the bottom. Liquid media (4 ml per flask), solid base (2 ml), and culture flasks were changed twice per day, and worms were washed three times in PBS at 37 °C with every change. For pulse and chase experiments with EdU, worms were incubated with 10 μM EdU for 2 h (pulse) and subsequently kept in EdU-free culture medium for 24, 48, and 96 h (chase) prior to fixation and downstream analysis.

## Acknowledgments

The authors gratefully acknowledge the Advanced Bioimaging Unit at the Institut Pasteur Montevideo for their support and assistance in the present work, and Beatriz Munguía, Laboratorio de Experimentación Animal, Facultad de Química, Universidad de la República, Uruguay for collaborating with the maintenance of the life cycle of *H. microstoma*.

---

**Supplementary Figure 1. Changes in gene expression associated with reproductive duct development in *H. microstoma*. A.** Expression of *hm-pc2* during the early, mid and late D4 stage of the development of the reproductive ducts of *H. microstoma.* **B.** Expression of *hm-fzd5/8* during the development of the reproductive ducts (stages D2 and D4). Bars: 50 μm. Abbreviations: def, vas deferens; ef, efferent ducts; esv, external seminal vesicle; isv, internal seminal vesicle; sr, seminal receptacle; t, testes; vag, vagina.

**Supplementary Figure 2.** Increase in the number of proglottids during reproductive development of *H. diminuta* starting from day 4 post-infection. The data was extracted from tables X, XI, and XII of Roberts et al., 1961.^45^

## Notes

**Grant support information:** this work was supported by Dirección para el Desarrollo de la Ciencia y el Conocimiento, Grant/Award Number: FVF2017/014, and Programa de Desarrollo de las Ciencias Básicas (no specific grant number applicable).

### Competing Interest Statement

The authors have declared no competing interest.

