## Supplementary figures and images for "Coordinated development of the male and female reproductive systems in the cestode *Hymenolepis microstoma*"

### Supplementary Figure 1

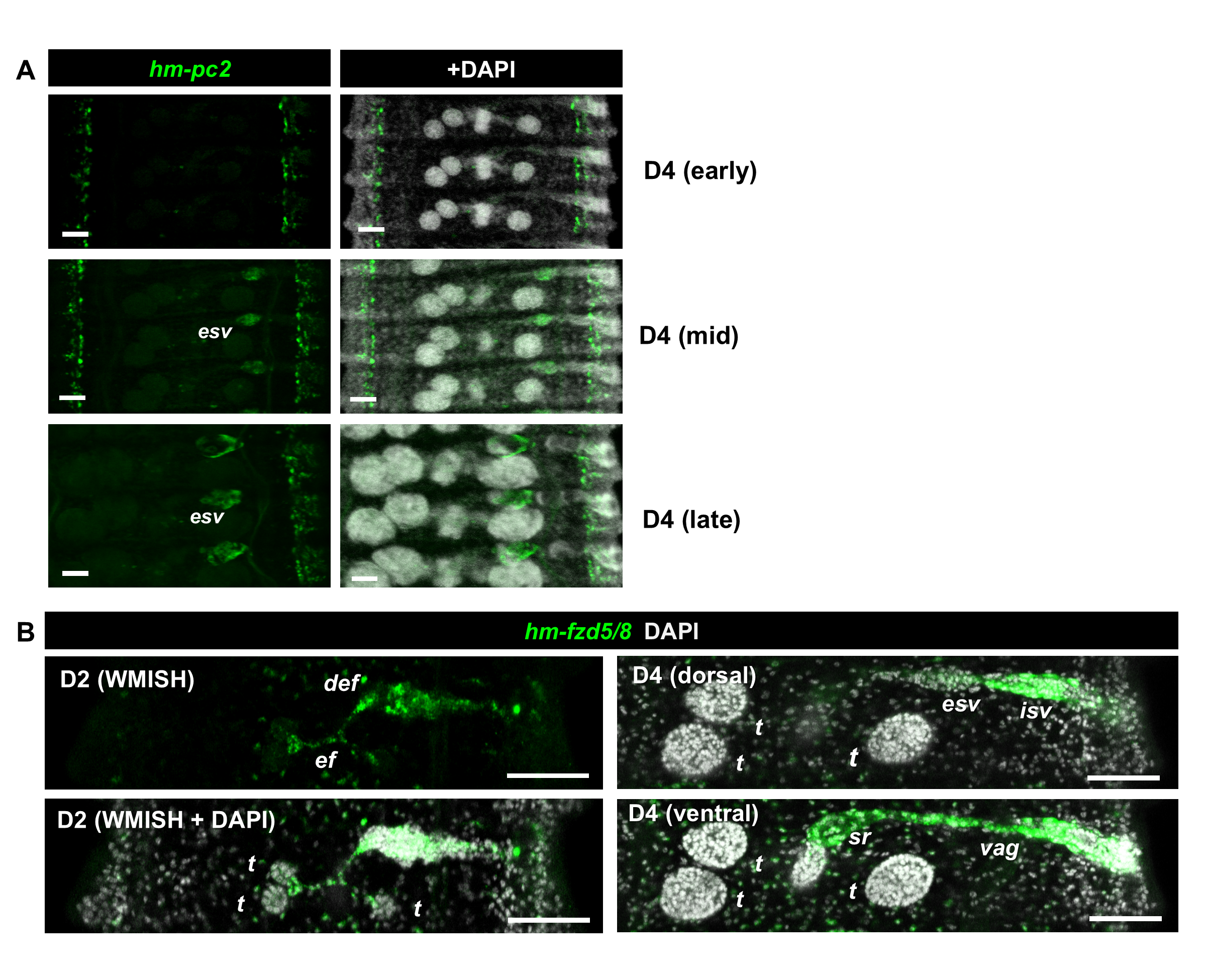

### Supplementary Figure 2

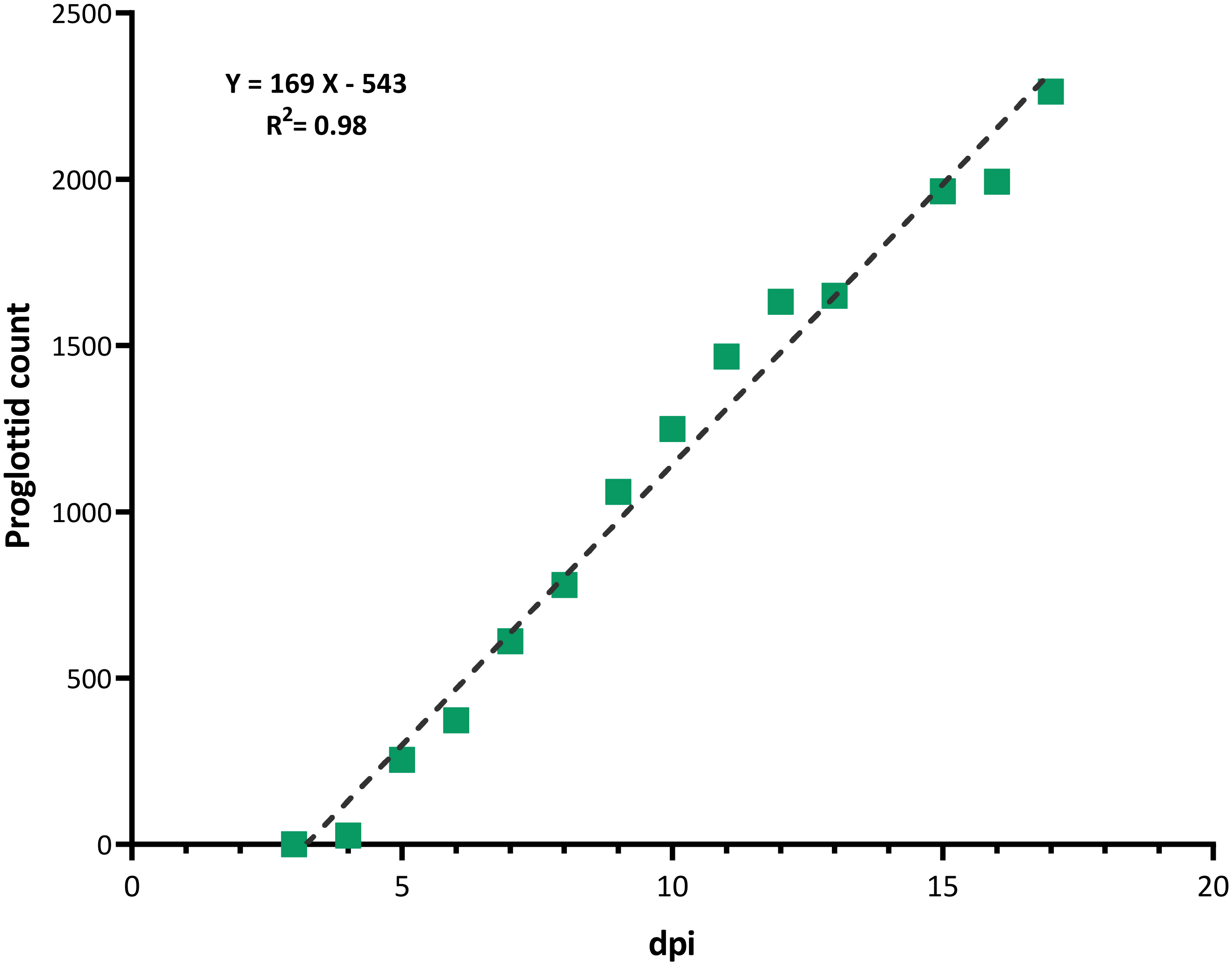
